# Emotion Recognition in a Multi-Componential Framework: The Role of Physiology

**DOI:** 10.1101/2021.04.08.438559

**Authors:** Maëlan Q. Menétrey, Gelareh Mohammadi, Joana Leitão, Patrik Vuilleumier

## Abstract

Emotions are rich and complex experiences involving various behavioral and physiological responses. While many empirical studies have focused on discrete and dimensional representations of emotions, these representations do not fully reconcile with recent neuroscience studies that increasingly suggest a multi-process mechanism underlying emotional experience. Moreover, the latter view accords with psychological theories that consider emotions as multicomponent phenomena, such as appraisal theories. Although there is no complete consensus on the specific components of emotions and fundamental principles defining their organization, the Component Process Model (CPM) is well established framework describing an emotion as a dynamic process with five major highly interrelated components: cognitive appraisal, expression, motivation, physiology and feeling. Yet, few studies have systematically investigated a range of discrete emotions through this full multi-componential view. In the present study, we therefore elicited various emotions during movie watching and measured their manifestation across these components. Our primary goal was to investigate the relationship between physiological measures and the theoretically defined components of emotions. In addition, we also investigated whether discrete emotions could be predicted from information provided by the multicomponent response patterns, as well as the specific contributions of each component in such predictions. Results suggest that physiological features are interrelated to all other components of emotion, but the least significant predictors for emotion classification. Overall, emotion prediction was significantly higher when classifiers were trained with all five components. The findings therefore support a description of emotion as a dynamic multicomponent process, in which the emergence of a conscious feeling state requires the integration of all the components.

## INTRODUCTION

Emotions play a central role in human experience by changing the way we think and behave. However, our understanding of the complex mechanisms underlying their production still remains incomplete and debated. Various theoretical models have been proposed to deconstruct emotional phenomena by highlighting their constituent features, as well as the particular behaviors and particular feelings associated with them. Despite ongoing disagreements, there is a consensus at least in defining an emotion as a multicomponent response, rather than a unitary entity (Moors, 2009). This conceptualization concerning the componential nature of emotion is not only central in appraisal theories (Scherer, 2019) and constructivist theories (Barrett et al., 2007), but also found to some extent in dimensional (Russell, 2009) and basic categorical models (Matsumoto & Ekman, 2009) that consider emotions as organized along orthogonal factors of “core affect” (valence and arousal), or as discrete and modular adaptive response patterns (fear, anger, etc.), respectively. Among these, appraisal theories, such as the Component Process Model (CPM) of emotion proposed by Scherer (1984), provide an explicit account of emotion elicitation in terms of a combination of a few distinct processes that evaluate the significance and context of the situation (e.g., relevance, novelty, controllability, etc.) and triggers a set of synchronized and interdependent responses at different functional levels in both the mind and body (Scherer, 2009). Hence, it is suggested that multiple and partly parallel appraisal processes operate to modify the motivational state and in turn modulate the autonomic system (leading to somatovisceral changes), the somatic system (producing motor expression in face or voice and bodily actions), as well as the cognitive system (including attention, memory, etc.). Synchronized changes in all these components - appraisal, action tendencies, physiology, motor expression, and cognition – may be centrally integrated in a multimodal representation that eventually becomes conscious and constitutes the subjective feeling component of the emotion (Grandjean et al., 2008).

Because the CPM proposes to define an emotion as a bounded episode characterized by a particular pattern of component synchronization, whereby the degree of coherence among components is a central property of emotional experience (Scherer, 2005a), it offers a valuable framework to model emotions in computationally tractable features. Yet, previous studies often relied on physiological changes combined with subjective feeling measures, either in the perspective of discrete emotion categories (e.g., fear, anger, joy, etc.) or more restricted dimensional descriptors (e.g., valence and arousal) (see Gunes & Pantic, 2010). As a consequence, such approaches have generally overlooked the full componential view of emotion. On the other hand, studies inspired by the appraisal framework have often analyzed emotional response with linear analyses and simple linear models (Fontaine et al., 2013; Frijda et al., 1989; Smith & Ellsworth, 1985). Yet, based on the interactional and multicomponent account of emotions in this framework (Sander et al., 2005), nonlinear classification techniques from the field of machine learning may be more appropriate and indeed provide better performances in the discrimination of emotions (Meuleman et al., 2019; Meuleman & Scherer, 2013). However, in the few studies using such approaches, classification analyses were derived from datasets depicting the semantic representation of major emotion words, but participants were not directly experiencing genuine emotions.

In parallel, while physiology is assumed to be one of the major components of emotion, the most appropriate channels of physiological activity to assess or to differentiate a particular emotion is still debated (see Harrison et al., 2013). For example, dimensional and constructivist theories do not assume that different emotions present specific patterns of physiological outputs (Quigley & Barrett, 2014) or argue that evidence is minimal for supporting specific profiles in each emotional category, spotlighting the insufficient consistency and specificity in patterns of activation within the peripheral and central nervous systems (Siegel et al., 2018; Wager et al., 2015). It has also been advocated that an emotion emerges from an ongoing constructive process that involves a set of basic affect dimensions and psychological components that are not specific to emotions (Barrett et al., 2007; Lindquist et al., 2013). Therefore, the modulation of autonomic nervous system (ANS) activity might be tailored to the specific demand of a situation and not to a discrete emotion. Peripheral physiological state occurring during a given emotion type is therefore expected to be highly variable in its physiological nature.

In contrast, some authors argue that measures of peripheral autonomic activity may contain diagnostic information enabling the representation of discrete emotions, that is, a shared pattern of bodily changes within the same category of emotion that becomes apparent only when considering a multidimensional configuration of simultaneous measures (Kragel & LaBar, 2013). Because univariate statistical approaches, which evaluate the relationship between a dependent variable and one or more experimental independent variables, have shown inconsistent results in relating physiology measures to discrete emotions (Kreibig, 2010), the development of multivariate statistical approaches to discriminate multidimensional patterns offers new perspectives to address these issues. By assessing the correlation between both dependent and independent variables and by jointly considering a set of multiple variables, multivariate analyses can reveal a finer organization in data as compared with univariate analyses where variables are treated independently. Accordingly, several recent studies used multivariate techniques and described separate affective states based on physiological measures including cardiovascular, respiratory and electrodermal activity (Christie & Friedman, 2004; Kreibig et al., 2007; Stephens et al., 2010). Such results support theoretical accounts from both basic (Ekman, 1992) and appraisal models (Scherer, 1984) suggesting that information carried in autonomic responses is useful to distinguish between emotional states. In this view, by using the relationships between multiple physiological responses in different emotional situations, it should be possible to infer which emotion is elicited. However, empirical evidence suggests that it is still complicated to figure out from patterned physiological responses, whether ANS measures are differentiated among specific emotion categories or more basic dimensions (Quigley & Barrett, 2014; Mauss & Robinson, 2009). Moreover, it is often observed that self-reports of emotional experience discriminate between discrete emotions with a much better accuracy than autonomic patterns (Mauss & Robinson, 2009).

In sum, there is still no unanimous conclusion about distinguishable patterns of activation in ANS, due to the difficulty to identify and associate reliable response patterns to discrete emotions. As a consequence, the debate is not closed concerning the functionality of physiology during an emotional experience. Therefore, to provide further insights about the contribution of physiology in emotion differentiation, we propose here to examine how a full componential model can account for the multiple and concomitant changes in physiological and behavioral measures observed during emotion elicitation. To the best of our knowledge, the present work represents one of the first attempts to investigate the componential theory by explicitly considering a combination of multiple, theoretically defined, emotional processes that occur in response to naturalistic emotional events (from cinematic film excerpts). By deploying a data-driven computational approach based on multivariate pattern classification, we aim at performing detailed analyses of physiological data in order to distinguish and predict the engagement of different emotion components across a wide range of eliciting events. On the grounds of such multicomponent response patterns, we also aim at determining to what extent discrete emotion categories can be predicted from information provided by these components, and what is the contribution of each component in such predictions. We hypothesize that a multicomponent account, as proposed by the CPM (Scherer, 1984; 2009), may allow us to capture the variability of physiological activity during emotional episodes, as well as their differentiation across major categories of emotions.

## MATERIAL AND METHODS

Assuming that a wide range of emotional sequences will engage a comprehensive range of component processes, we selected a number of highly emotional film excerpts taken from different sources (see below). Physiological measures were recorded simultaneously during the initial viewing of movie clips, with no instructions other than be spontaneously absorbed by the movies. Participants were asked, during a second presentation, to fill out a detailed questionnaire with various key descriptors of emotion-eliciting episodes derived from the componential model (i.e., CoreGRID items) that assess several dimensions of appraisal, motivation, expression, physiology, and feeling experiences (Fontaine et al., 2013). We then examined whether the differential patterns of physiological measures observed across episodes could be linked to a corresponding distribution of ratings along the CoreGRID items, and whether the combined assessment of these items and physiological measures could be used together to distinguish between discrete emotions.

### Population

A total of 20 French-speaking and right-handed students (9 women, 11 men) between 19 and 25 years old (mean age = 20.95, SD = 1.79) took part in the main study. All of them reported no history of neurological or psychiatric disorder, gave a written informed consent after a full explanation of the study and were remunerated. This work was approved by Geneva Cantonal Research Committee and followed their guidelines in accordance with Helsinki declaration.

### Stimuli selection

To select a set of emotionally engaging film excerpts which could induce variations along different dimensions of the component model, a first preliminary study was conducted in separate study (Mohammadi et al., 2019; Mohammadi & Vuilleumier, 2020). We selected a set of 139 film clips from the previous literature on emotion elicitation, matching in terms of time and visual quality (Gross & Levenson, 1995; Philippot, 1993; Schaefer et al., 2010). Emotion assessment was collected in terms of discrete emotion labels and componential model descriptors. Initially, clips were evaluated over 14 discrete emotions (fear, anxiety, anger, shame, warm-hearted, joy, sadness, satisfaction, surprise, love, guilt, disgust, contempt, calm) based on a modified version of the Differential Emotion Scale (Izard et al., 1993; McHugo et al., 1982). For the component model, 39 descriptive items were selected from the CoreGRID instrument, capturing emotion features along the 5 components of interest: appraisal, motivation, expression, physiology, and feeling (Fontaine et al., 2013). This selection was performed based on the applicability to emotion elicitation scenarios while watching an event in a clip. The study was performed on Crowdflower, a crowdsourcing platform, and a total number of 638 workers participated. Based on average ratings and discreteness, 40 film clips were selected for this study (for more details, see Mohammadi & Vuilleumier, 2020). Shame, warm-hearted, guilt and contempt were excluded from the list of elicited emotions because no clips received high ratings for these four emotions.

Finally, another preliminary study was conducted to isolate the highest emotional moments in each clip. To this aim, 5 different participants watched the full clips and rated the emotional intensity of the scene using CARMA, a software for continuous affect rating and media annotation (Girard, 2014). The five annotations were integrated to find the most intense emotional events in each time series.

The final dataset was thus represented by 4 clips for each of the 10 selected discrete emotions, with a total duration of 74 minutes (average length of 111 seconds per clip). Moreover, between 1 to 4 highly emotional segments of 12 seconds were selected in each film excerpt, for a total of 119 emotional segments. The list of the 40 selected films in our final dataset is presented in Supplementary Material (Table S1). For each film excerpt, the duration, the assigned emotion label and the number of highly emotional segments are indicated.

### Experimental paradigm

The whole experiment consisted of 4 sessions scheduled on different days. Each session was divided into two parts, fMRI experiment and behavioral experiment, lasting for about 1h and 2h, respectively. In the current study we focus only on the behavioral analysis and will not use the fMRI data. Stimuli presentation and assessment were controlled using Psychtoolbox-3, an interface between MATLAB and computer hardware.

During the fMRI experiment, participants were engaged in an emotion elicitation procedure using our 40 emotional film excerpts. No explicit task was required during this phase. They were simply instructed to let themselves feel and express emotions freely rather than controlling feelings and thoughts because of the experiment environment. Movies were presented inside the MRI scanner on an LCD screen through a mirror mounted on the head coil. The audio stream was transmitted through MRI-compatible earphones. Each session was composed of 10 separate runs, each presenting a film clip preceded by a 5-seconds instruction screen warning about the imminent next display and followed by a 30-seconds washout periods introduced as a low-level perceptual control baseline for the fMRI analysis (not analysed here). Moreover, a session consisted of a pseudo-random choice of 10 unique film clips with high ratings on at least one of the 10 different pre-labeled discrete emotion categories (fear, anxiety, anger, joy, sadness, satisfaction, surprise, love, disgust, calm). This permitted to engage potentially different component processes in every session. To avoid any order effect, the presentation of all stimuli was counterbalanced.

The behavioral experiment was performed at the end of each fMRI session, in a separate room. Participants were let alone with no imposed time constraints to complete the assessment. They were asked to rate their feelings, thoughts, or emotions evoked during the first viewing of the film clips and advised not to report what might be expected to feel in general when watching such kinds of events. To achieve the emotion evaluation, the 10 film excerpts seen in the preceding session were presented on a laptop computer with LCD screen and headphones. However, the previously selected highly emotional segments (see “stimuli selection” above) were now explicitly highlighted in each film excerpts by a red frame surrounding the visual display. In order to ensure that emotion assessment corresponded to a single event and not the entire clip, the ratings were required right after each segment by pausing the clips. The assessment involved a subset of CoreGRID instrument (Fontaine et al., 2013), which is to date the most comprehensive attempt for multi-componential measurement in emotion. The set of 32 items (see Table 1) had been pre-selected based on their applicability to the emotion elicitation scenario with movies, rather than according to an active first-person involvement in an event. Among our set of CoreGRID items, 9 were related to the appraisal component, 6 to the expression component, 7 to the action tendencies component, 6 to the feeling component, and 4 to the bodily component. Participants had to indicate how much they considered that the description of the CoreGRID items correctly represented what they felt in response to the highlighted segment, using a 7-level Likert scale with 1 for “not at all” and 7 for “strongly”. All responses were collected through the keyboard, for a total of 3808 assessments per participant (32 items × 119 emotional segments). Finally, they were also asked to label the segments by selecting one discrete emotion term from the list of 10 emotion categories. The frequency histogram showing the categorical emotions selected by the participants is presented in Supplementary Material section (Figure S1)

**Table 1.** List of the 32 CoreGRID items. Participants were asked to indicate on a 7-point Likert scale how much the descriptions represented what they felt.

| Major Components | CoreGrid items |
| --- | --- |
| **Appraisal** |  |
| To what extent did you ... | 1) think it was incongruent with your standards and ideas?<br>2) feel it was unpleasant for you?<br>3) think it violated laws or socially accepted norms?<br>4) think it was unpleasant for somebody (in the clip)?<br>5) think it was important and relevant for the goals or needs of somebody?<br>6) feel the event was unpredictable?<br>7) feel the event occurred suddenly?<br>8) think the event was caused by chance?<br>9) think the consequences were predictable? |
| **Expression** |  |
| To what extent did you ... | 10) press lips together?<br>11) close your eyes?<br>12) show tears?<br>13) have the jaw drop?<br>14) have eyebrows go up?<br>15) produce abrupt body movements? |
| **Action** |  |
| To what extent did you ... | 16) want to destroy something?<br>17) want to do damage, hit or say something that hurts?<br>18) feel the urge to stop what was happening?<br>19) want to undo what was happening?<br>20) want the ongoing situation to last or be repeated?<br>21) feel motivated to pay attention to what was going on?<br>22) want to tackle the situation and do something? |
| **Feeling** |  |
| To what extent did you ... | 23) feel bad?<br>24) feel calm?<br>25) feel good?<br>26) feel strong?<br>27) feel an intense emotional state?<br>28) experience an emotional state for a long time? |
| **Body** |  |
| To what extent did you ... | 29) experience muscles tensing (whole body)?<br>30) have a feeling of a lump in the throat?<br>31) have stomach troubles?<br>32) feel warm? |

### Physiological data acquisition

A number of physiological measures were collected during the first part of each session in the MRI scanner, including heart rate, respiration rate, and electrodermal activity. All the measures were acquired continuously throughout the whole scanning time. The data were first recorded with a 5000Hz sampling rate using the MP150 Biopac Systems software (Santa Barbara, CA), before being pre-processed with AcqKnowledge 4.2 and MATLAB 2012b.

Heart rate (HR) was recorded with a photoplethysmogram amplifier module (PPG100C). This single channel amplifier designed for indirect measurement of blood pressure was coupled to a TSD200-MRI photoplethysmogram transducer fixed on the index finger of the left hand. Recording artifacts and signal losses were corrected using endpoint function from AcqKnowledge, which interpolates the values of a selected impaired measure portion. Secondly, the pulse signal was exported to MATLAB and downsampled to 120 Hz. To remove scanner artefacts, a comb-pass filter was applied at 17,5 Hz. The pulse signal was then filtered with a band-pass filter between 1 and 40 Hz. Subsequently, the instantaneous heart rate was computed by identifying the peaks in the pulse signal, calculating the time intervals between them and converting this distance into beats per minute (BPM). The standard heart rate in humans goes from 60-100 bpm at rest. Hence, it was considered that a rate above 100 bpm was unlikely and the minimum distance between peaks will not exceed this limit. This automatic identification was manually verified by adding, changing or removing the detected peaks and possible outliers.

Respiration rate (RR) was measured using a RSP100C respiration pneumogram amplifier module, designed specifically for recording respiration effort. This differential amplifier worked with a TSD201 respiration transducer, which was attached with a belt around the upper chest near the level of maximum amplitude in order to measure thoracic expansion and contraction. Using a similar procedure as for HR preprocessing, the connect endpoint function of AcqKnowledge was first employed to correct manually the artifacts and losses of signal. After exporting the raw signal to MATLAB, it was downsampled to 120 Hz and then filtered with a band pass filter fixed between 0.05 and 1 Hz. Lastly, the signal was converted to breaths per minute using the same procedure as above. The standard respiration rate in human goes from 12 to 20 breaths per minute at rest. Since participants were performing a task inside a scanner which could be an unusual environment, the higher maximum rate was increased at 35 cycles per minute. Therefore, it was estimated that a rate above 35 was unlikely and the minimum distance between peaks will not exceed this limit. Again, this information was used in the automatic detection of the signal peaks. The respiration rate was then manually verified by looking at the detected signal peaks and corrected, with outliers being removed when it was necessary.

Electrodermal activity (EDA) was registered using an EDA100C electrodermal activity amplifier module, a single-channel, high-gain, differential amplifier designed to measure skin conductance via the constant voltage technique. The EDA100C was connected to Adult ECG Cleartrace 2 LT electrodes with conductive adhesive gel, which were placed on the index and the median fingers of the participants. Following the manual correction of artifacts and losses of signal with the connect endpoint function on AcqKnowledge, the raw signal was exported to MATLAB. Similar to the two other physiological signals, the EDA signal was downsampled to 120 Hz. This signal, recorded by BIOPAC in microSiemes (μS), was then filtered with a 1Hz low pass filter. An IIR (infinite impulse response) high-pass filter fixed at 0.05 Hz was applied to derive the Skin Conductance Responses (phasic component of EDA) representing the rapidly changing peaks, while a FIR (finite impulse response) low-pass filter fixed at 0.05Hz was applied to derive the Skin Conductance Levels (tonic component of EDA) corresponding to the smooth underlying slowly-changing levels (AcqKnowledge 4 Software Guide, 2011).

### Features and normalization

MATLAB was used to select physiological values during the 12-seconds duration of high emotional segments. From these values, the means, variances, and ranges of each physiological signals (HR, RR, phasic and tonic EDA) were calculated. For HR and RR measures, respectively 4 and 17 responses during highly emotional segments had to be removed in one participant due to a corrupted signal, potentially induced by movement. For EDA, 267 values had to be removed due to temporary losses of signal, resulting in flat and useless measures. In particular, EDA responses of two subjects were completely removed as the EDA sensor could not capture their response. In order to handle the missing values in the physiological data, dropouts were replaced by mean value of the whole session during which the signal loss has happened. Furthermore, the variance in physiological responses could be very large and different across participants. Because it was particularly important to reduce such variability in order to avoid inter-individual biases, all physiological measures were normalized within-subject using RStudio (1.1.383). To achieve this, standardized z-scores were calculated from the physiological data during the 4 sessions of each participant.

Regarding responses collected for the 32 CoreGRID items for each high emotional segment, a within-subject normalization into z-score was also performed. These behavioral data and the discrete emotion labels selected by the participants for each emotional segment were entered in our analysis to determine their association with the related physiological measures.

### Predictive analyses

To investigate the relationship between physiology and the component model descriptors, two analyses were performed. First, we examined whether the physiology measures allowed predicting component model descriptors and vice versa. Second, we assessed whether distinct features from the componential model allowed predicting discrete emotion categories and compared the value of different components for this prediction. For both analyses, multivariate pattern classifications using machine learning algorithms were undertaken to predict the variables of interest. Linear and nonlinear classifiers including Logistic Regression (LR) and Support Vector Machine (SVM) with different kernels (linear, radial basis function, polynomial and sigmoid) were applied. All analyses were carried out using the RStudio statistical software, Version 1.1.383. The binary and multiclass classifications using Support Vector Machine were conducted with the “e1071” package, Version 1.7.

In order to permit such analyses and to simplify the computational problem, the scores of the dependent variable were converted into two classes of “High” and “Low” using the median value across all the participants as a cutoff threshold. First, the CoreGRID items were used as predictor variables to predict the dependent variable, which was either the mean or the variance of each physiological measure. In particular, LR and SVM with linear and non-linear kernels using 10-fold cross-validation were applied. To guarantee test and training independence, each participant’s assessment was included in either a test set or a training set. Second, similar analyses were carried out to determine whether physiology measures could encode the component model descriptors, but now using the physiology measures as independent variables in an attempt to predict the ratings of each CoreGRID item as either above or below the median. For both analyses, SVM with radial basis function (RBF) kernel outperformed LR and other SVM models in almost all cases, so we will only report the result from SVM with RBF kernel. Finally, to examine the relationship between the component process model and discrete emotion types, multiclass classifications using SVM with RBF kernel were performed on different combinations of CoreGRID items and physiological measures in order to predict specific emotion categories as labeled by the participants. Given the large number of classes and limited number of samples per class with too many predictors, a leave-one-subject-out cross-validation was used to guarantee a complete independence between the training and testing datasets.

## RESULTS

### Multivariate pattern analyses: Binary classifications

Our predictive analyses aimed to assess classification based on multivariate patterns using either physiology measures or CoreGRID items for different emotional movie segments, treated as binary dependent variables (High vs Low).

In the first instance, we deployed SVM classifications with 10-fold cross-validation to predict each physiological measure (mean or variance) as high or low from responses to the 32 CoreGRID items. This classifier yielded accuracies significantly greater than the chance rate of 50% for all physiological measures (Table 2). However, while these binary classifications were statistically significant and effect sizes were large, their discriminative performance remained weak (on average, about 58% of correct responses).

**Table 2.** Cross-subject SVM classifications. Predictions of physiological changes from the 32 CoreGRID items. Accuracy rate represents the percentage of correct classifications. Paired t-tests were conducted to verify significant differences between SVM classifier and chance level. As estimates of effect size, we report Cohen’s d and 95% confidence interval.

| Binary classification |  | Accuracy | Chance level | t | df | p-value | Cohen's D | 95% CI |
| --- | --- | --- | --- | --- | --- | --- | --- | --- |
| **SVM classifier** |  |  |  |  |  |  |  |  |
| HR | mean | 0.59 | 0.5 | 7.175 | 9 | < **0.001** | 3.497 | [0.76 6.23] |
|  | variance | 0.55 | 0.5 | 4.308 | 9 | **0.002** | 1.480 | [0.43 2.52] |
| RR | mean | 0.56 | 0.5 | 8.490 | 9 | < **0.001** | 2.950 | [1.26 4.63] |
|  | variance | 0.54 | 0.5 | 7.133 | 9 | < **0.001** | 2.246 | [1.00 3.48] |
| phasic EDA | mean | 0.58 | 0.5 | 8.549 | 9 | < **0.001** | 4.494 | [0.81 8.17] |
|  | variance | 0.61 | 0.5 | 12.641 | 9 | < **0.001** | 2.616 | [1.70 3.53] |
| tonic EDA | mean | 0.59 | 0.5 | 10.691 | 9 | < **0.001** | 5.231 | [1.29 9.17] |
|  | variance | 0.60 | 0.5 | 17.761 | 9 | < **0.001** | 7.062 | [2.08 11.31] |

Conversely, using the same classification approach, the ratings of each CoreGRID item were predicted from the combination of physiological measures and physiology items in the CoreGRID questionnaire. Results from SVM showed that a majority of the CoreGRID items could be predicted significantly better than the chance level (Table 3), however, the classification accuracies were still relatively low (on average, about 55% of correct responses). The most reliable discrimination levels (highest t values relative to chance rate) were observed for appraisals and feelings of unpleasantness (55% of correct responses) as well as action tendencies (want to destroy / to do damage, 58% of correct responses).

**Table 3.**
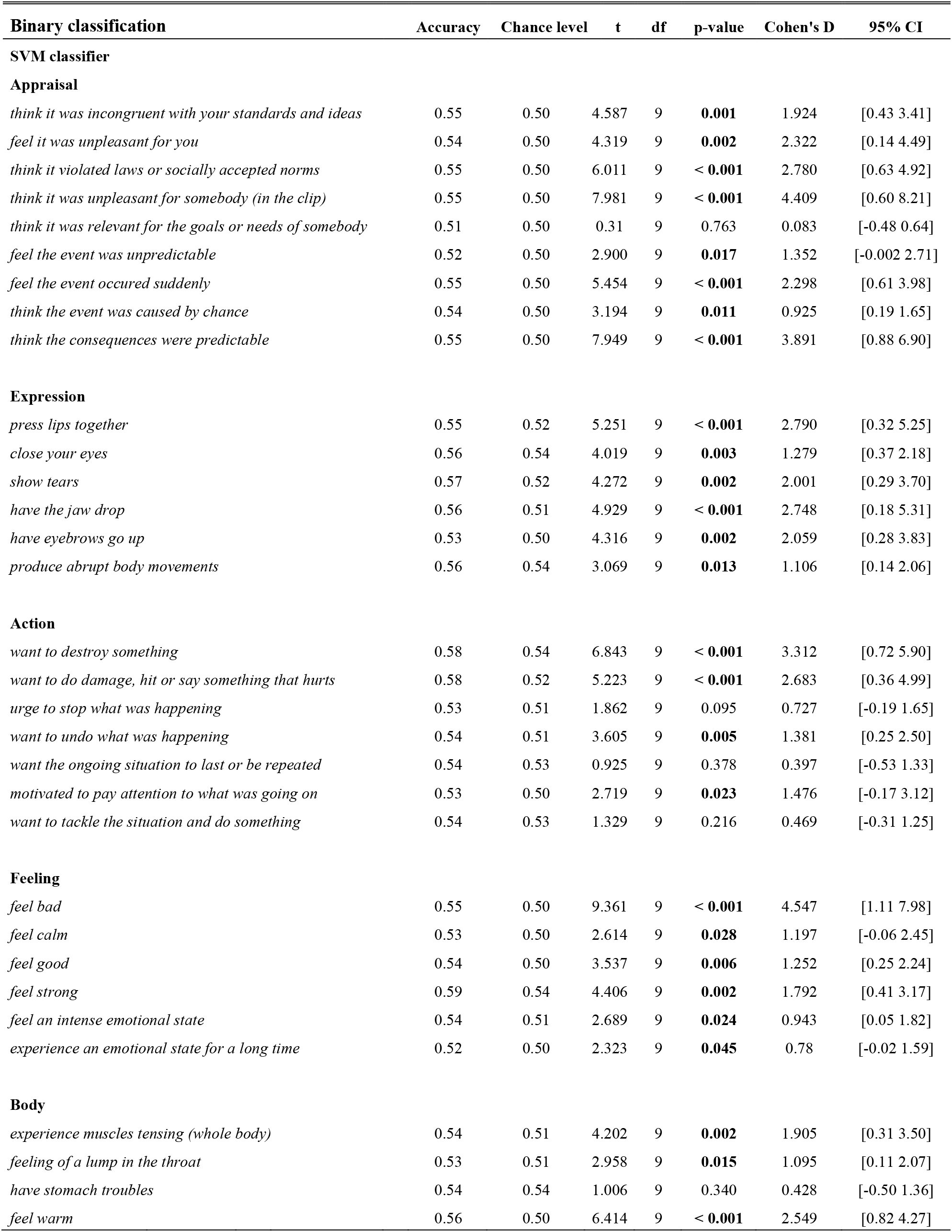
Cross-subject SVM classifications. Predictions of individual CoreGRID item ratings from the mean and variance measures of physiological responses. Accuracy rate represents the percentage of correct classifications. Paired t-tests were conducted to test for significant differences between SVM classifier and chance level. As estimates of effect size, we report Cohen’s d and 95% confidence interval.

| Binary classification | Accuracy | Chance level | t | df | p-value | Cohen's D | 95% CI |
| --- | --- | --- | --- | --- | --- | --- | --- |
| **SVM classifier** |  |  |  |  |  |  |  |
| **Appraisal** |  |  |  |  |  |  |  |
| *think it was incongruent with your standards and ideas* | 0.55 | 0.50 | 4.587 | 9 | **0.001** | 1.924 | [0.43 3.41] |
| *feel it was unpleasant for you* | 0.54 | 0.50 | 4.319 | 9 | **0.002** | 2.322 | [0.14 4.49] |
| *think it violated laws or socially accepted norms* | 0.55 | 0.50 | 6.011 | 9 | **< 0.001** | 2.780 | [0.63 4.92] |
| *think it was unpleasant for somebody (in the clip)* | 0.55 | 0.50 | 7.981 | 9 | **< 0.001** | 4.409 | [0.60 8.21] |
| *think it was relevant for the goals or needs of somebody* | 0.51 | 0.50 | 0.31 | 9 | 0.763 | 0.083 | [-0.48 0.64] |
| *feel the event was unpredictable* | 0.52 | 0.50 | 2.900 | 9 | **0.017** | 1.352 | [-0.002 2.71] |
| *feel the event occurred suddenly* | 0.55 | 0.50 | 5.454 | 9 | **< 0.001** | 2.298 | [0.61 3.98] |
| *think the event was caused by chance* | 0.54 | 0.50 | 3.194 | 9 | **0.011** | 0.925 | [0.19 1.65] |
| *think the consequences were predictable* | 0.55 | 0.50 | 7.949 | 9 | **< 0.001** | 3.891 | [0.88 6.90] |
| **Expression** |  |  |  |  |  |  |  |
| *press lips together* | 0.55 | 0.52 | 5.251 | 9 | **< 0.001** | 2.790 | [0.32 5.25] |
| *close your eyes* | 0.56 | 0.54 | 4.019 | 9 | **0.003** | 1.279 | [0.37 2.18] |
| *show tears* | 0.57 | 0.52 | 4.272 | 9 | **0.002** | 2.001 | [0.29 3.70] |
| *have the jaw drop* | 0.56 | 0.51 | 4.929 | 9 | **< 0.001** | 2.748 | [0.18 5.31] |
| *have eyebrows go up* | 0.53 | 0.50 | 4.316 | 9 | **0.002** | 2.059 | [0.28 3.83] |
| *produce abrupt body movements* | 0.56 | 0.54 | 3.069 | 9 | **0.013** | 1.106 | [0.14 2.06] |
| **Action** |  |  |  |  |  |  |  |
| *want to destroy something* | 0.58 | 0.54 | 6.843 | 9 | **< 0.001** | 3.312 | [0.72 5.90] |
| *want to do damage, hit or say something that hurts* | 0.58 | 0.52 | 5.223 | 9 | **< 0.001** | 2.683 | [0.36 4.99] |
| *urge to stop what was happening* | 0.53 | 0.51 | 1.862 | 9 | 0.095 | 0.727 | [-0.19 1.65] |
| *want to undo what was happening* | 0.54 | 0.51 | 3.605 | 9 | **0.005** | 1.381 | [0.25 2.50] |
| *want the ongoing situation to last or be repeated* | 0.54 | 0.53 | 0.925 | 9 | 0.378 | 0.397 | [-0.53 1.33] |
| *motivated to pay attention to what was going on* | 0.53 | 0.50 | 2.719 | 9 | **0.023** | 1.476 | [-0.17 3.12] |
| *want to tackle the situation and do something* | 0.54 | 0.53 | 1.329 | 9 | 0.216 | 0.469 | [-0.31 1.25] |
| **Feeling** |  |  |  |  |  |  |  |
| *feel bad* | 0.55 | 0.50 | 9.361 | 9 | **< 0.001** | 4.547 | [1.11 7.98] |
| *feel calm* | 0.53 | 0.50 | 2.614 | 9 | **0.028** | 1.197 | [-0.06 2.45] |
| *feel good* | 0.54 | 0.50 | 3.537 | 9 | **0.006** | 1.252 | [0.25 2.24] |
| *feel strong* | 0.59 | 0.54 | 4.406 | 9 | **0.002** | 1.792 | [0.41 3.17] |
| *feel an intense emotional state* | 0.54 | 0.51 | 2.689 | 9 | **0.024** | 0.943 | [0.05 1.82] |
| *experience an emotional state for a long time* | 0.52 | 0.50 | 2.323 | 9 | **0.045** | 0.78 | [-0.02 1.59] |
| **Body** |  |  |  |  |  |  |  |
| *experience muscles tensing (whole body)* | 0.54 | 0.51 | 4.202 | 9 | **0.002** | 1.905 | [0.31 3.50] |
| *feeling of a lump in the throat* | 0.53 | 0.51 | 2.958 | 9 | **0.015** | 1.095 | [0.11 2.07] |
| *have stomach troubles* | 0.54 | 0.54 | 1.006 | 9 | 0.340 | 0.428 | [-0.50 1.36] |
| *feel warm* | 0.56 | 0.50 | 6.414 | 9 | **< 0.001** | 2.549 | [0.82 4.27] |

### Multivariate pattern analyses: Multiclass classifications

Our second aim was to investigate whether discrete emotion categories as indicated by the participants can be predicted from ratings of their componential profile. This more global pattern analysis for a multiclass variable requires to go further than simple binary classification. To achieve this, a multiclass SVM classifications with leave-one-subject-out cross-validation was performed, taking the combination of within-subject normalized mean and variance measures from the 4 physiological signals and all behavioral responses to the CoreGRID items for the 5 emotional components as predictors. Applying the SVM trained classifiers on separate testing datasets, we obtained an average accuracy rate of 45.4% in comparison to a rate of 17.6% for the chance level (*t*(19)= 10.852, *p* < 0.001, Cohen’s *d* = 3.41, 95% *CI* [1.75 5.06]) (Figure 1A).

**Figure 1.**
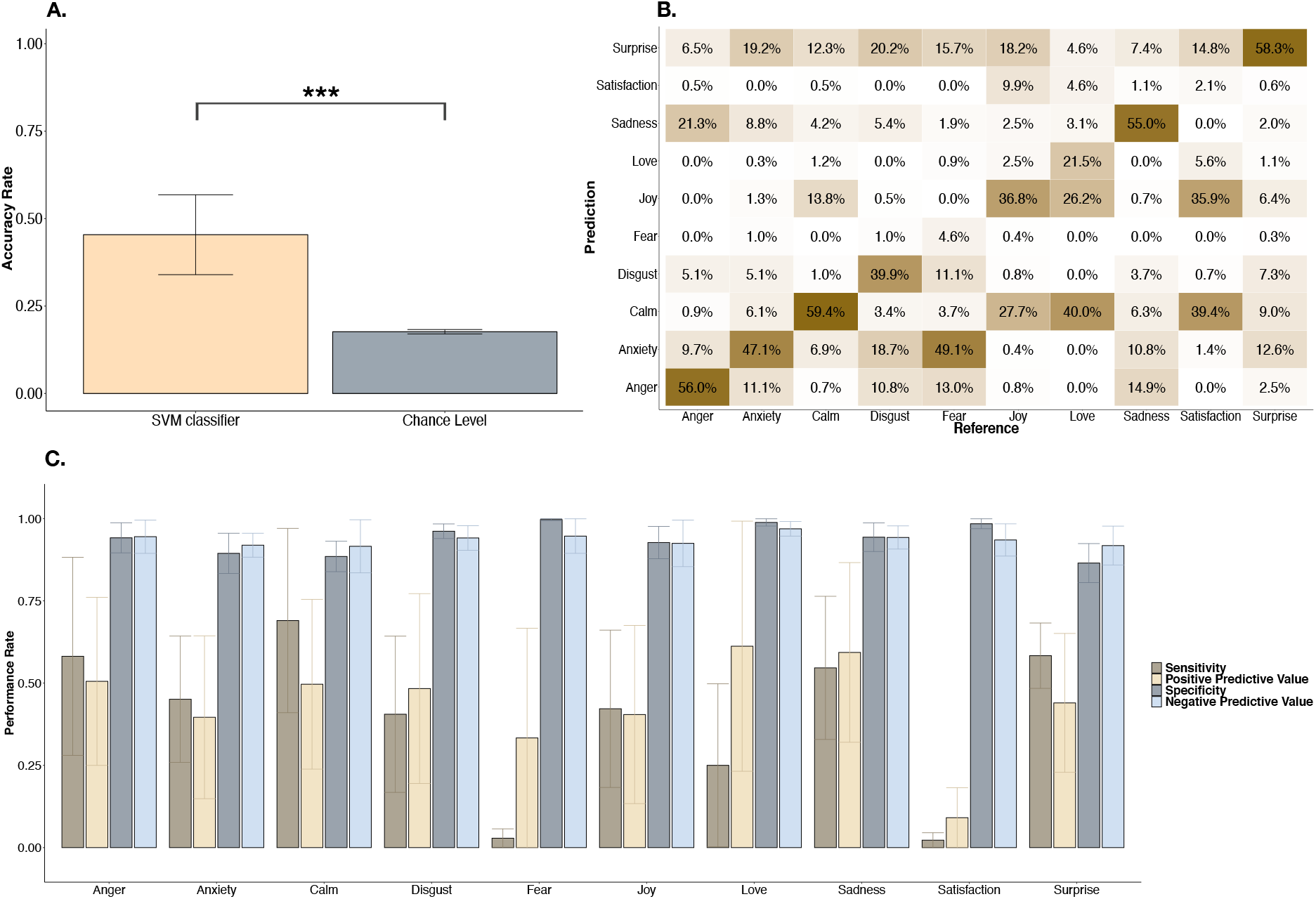
**A. Cross-subject multiclass SVM classification with leave-one-subject-out cross-validation**. Predictions of discrete emotion labels from the mean and variance of physiological measures and behavioral responses to the 32 CoreGRID features. Accuracy rate represents the percentage of correct classifications. The error bars show the standard deviation. Paired t-tests were conducted to assess significant differences between SVM classifier and chance level, as highlighted by asterisks indicating the p-value (*** *p* < 0.001). **B. Confusion matrix of emotion labels**. The diagonal running from the lower left to the upper right represents the correct predicted emotion. **C. Statistical measures of classification performances across emotions.** Average of statistical measures assessing the performances from the 20 classifiers. The error bars show the standard deviation.

The confusion matrix showed that five emotion categories (anger, calm, sadness and surprise) were correctly predicted more than half of the times, with an accuracy range from 55% to 59.4% (Figure 1B). By contrast, predictions were extremely unsuccessful for fear (misclassified as anxiety) and satisfaction (misclassified as joy or calm). However, it is worth noticing that these categorical emotions had a smaller number of instances since they were less often selected by the participants (see Figure S1). Interestingly, incorrect predictions for these two emotions were still related to some extent to the target category. Indeed, mainly anxiety but also disgust and surprise were predicted instead of fear, whereas joy and calm were predicted instead of satisfaction. Love was also frequently misclassified as joy and calm.

Concomitantly, statistical measures allowing the assessment of prediction performance indicated that specificity and negative predictive value were particularly high for all emotions (Figure 1C). This suggests that the classification algorithm had a notable ability to correctly reject observations that did not belong to the emotion of interest, that is, to provide a good degree of certainty and reliability for true negatives. In contrast, sensitivity and positive predictive value were not as good and fluctuated substantially across the emotions, with the best performance for calm and the worst for fear and satisfaction (Figure 1C).

Finally, to better identify the relation between discrete emotions and the different component processes, five multiclass SVM classifications with leave-one-subject-out cross-validation were performed using different combinations of these components. In every model, one component was excluded to investigate the added information brought by that component (Figure 2). In comparison to the accuracy rate of the complete model using physiology and all the CoreGRID items (45.4%), the accuracy rate of the reduced model without the body physiology component (4 CoreGRID items and all physiological measures) did not significantly change (45.2%, *t*(19)= −0.17, *p* = 0.866, Cohen’s *d* = −0.01, 95% *CI* [−0.13 0.11]). These results point to the fact that information coming from features of the body physiology component in our study was already encoded in other components. On the other hand, with respect to performance with the full model, reduced models without the appraisal component (41.4%, *t*(19)= −4.598, *p* < 0.001, Cohen’s *d* = −0.33, 95% *CI* [−0.48 −0.18]), without the expression component (41.9%, *t*(19)= −4.358, *p* < 0.001, Cohen’s *d* = −0.29, 95% *CI* [−0.43 −0.15]), without the motivation component (43%, *t*(19)= −2.489, *p* = 0.022, Cohen’s *d* = −0.21, 95% *CI* [−0.37 −0.03]) or without the feeling component (44%, *t*(19)= −2.349, *p* = 0.029, Cohen’s *d* = −0.12, 95% *CI* [−0.22 −0.02]) were statistically less predictive, even though the effects sizes remained relatively small.

**Figure 2.**
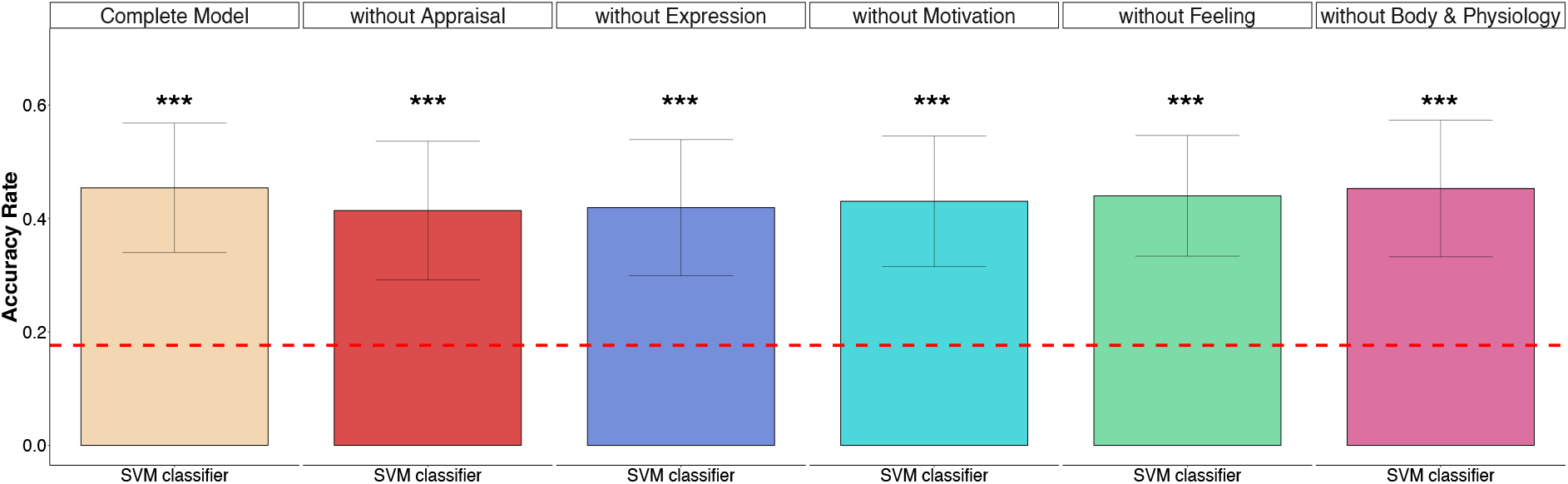
Emotion classification using one-component-out models. Cross-subject multiclass SVM classifications with leave-one-subject-out cross-validation. Predictions of discrete emotion categories from emotional components based on their related CoreGRID items, using one-component-out models, for each of the 5 components. Accuracy rate represents the percentage of correct classifications. The error bars show the standard deviation. Paired t-tests have been conducted and significant differences between SVM classifier and chance level are highlighted (asterisks indicate the p-value: *** *p* < 0.001).

In addition, the contributions of major components of emotion were assessed by predicting discrete emotion labels from each component separately. Five multiclass SVM classifications with leave-one-subject-out cross-validation were performed from the appraisal, expression, motivation, feeling, and body components (body items and all physiological measures). Since prediction performance from each of the 5 emotion components yielded accuracies significantly greater than the chance rate of 17.6% (Figure 3), we also analyzed the average sensitivity rate across the different classifiers in order to determine more precisely the power of each component to distinguish the different discrete emotions.

**Figure 3.**
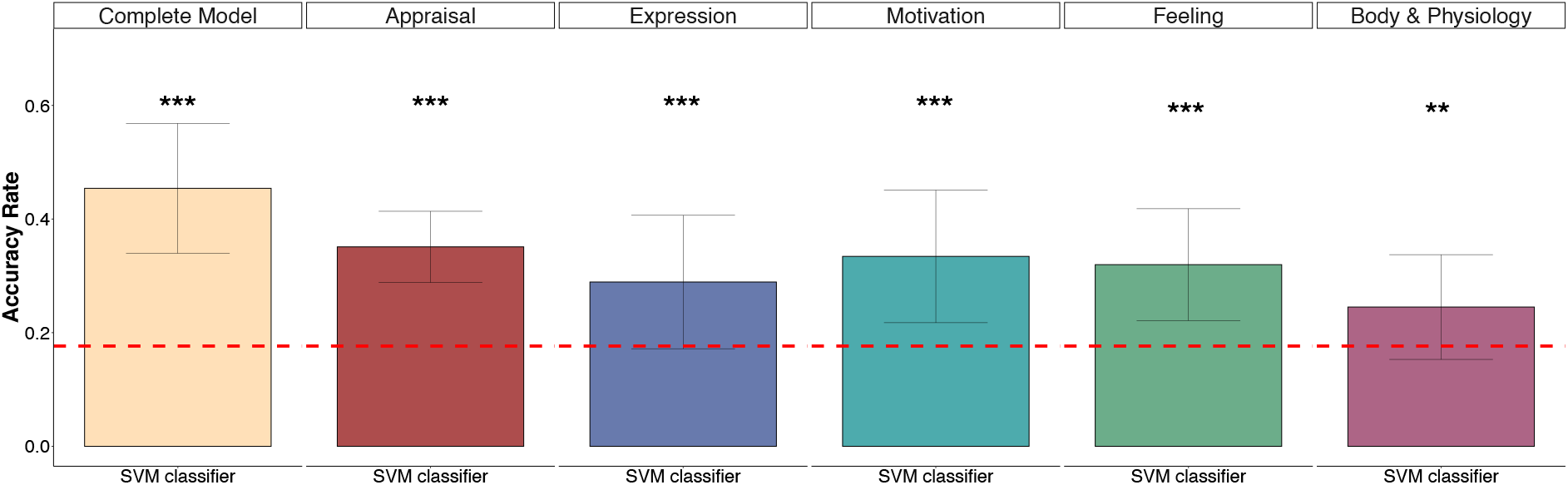
Emotion classification using the components independently. Cross-subject multiclass SVM classifications with leave-one-subject-out cross-validation. Predictions of discrete emotion labels from each component individually. Accuracy rate represents the percentage of correct classifications. The error bars show the standard deviation. Paired t-tests have been conducted and significant differences between SVM classifier and chance level are highlighted (asterisks indicate the p-value: ** *p* < 0.01, *** *p* < 0.001).

While the results above suggested that the percentage of correct predictions was generally similar regardless of the particular component used to train the classifiers, these additional analyses indicate that the pattern of features from specific components may yield a more reliable detection of particular emotions relative to others (Figure 4A). Moreover, it appeared also that some emotion classes were consistently well discriminated by all components (e.g., calm and sadness), while others (fear, love, and satisfaction) were poorly predicted by any component. Conversely, some components could have more importance for particular emotions (e.g., surprise is well predicted with appraisal but not body features, while motivation component features seem the best to predict anger and joy).

**Figure 4.**
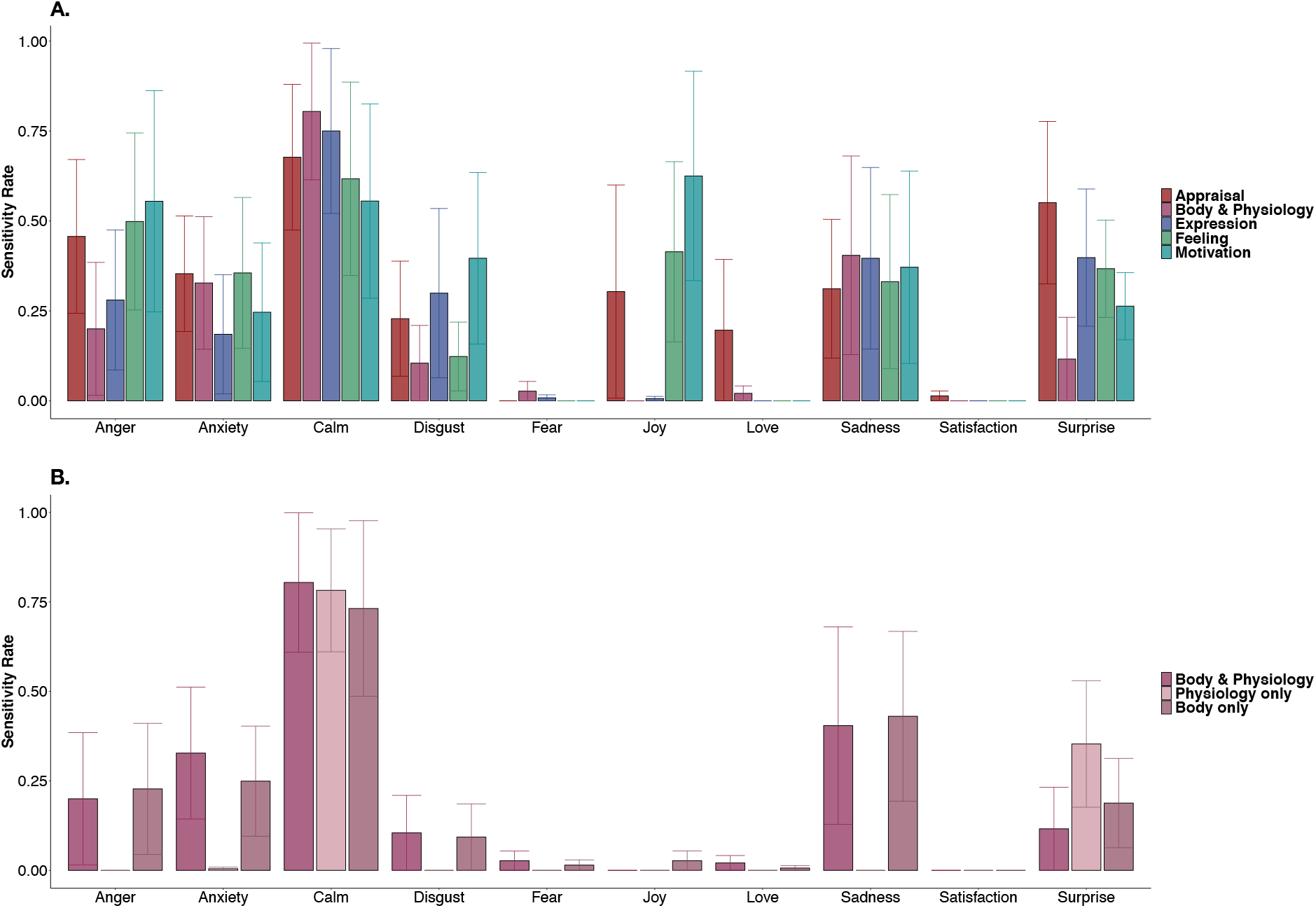
**A. Sensitivity of emotion detection from components assessed by the CoreGRID**. Prediction of discrete emotion labels from components assessed by the CoreGRID. **B. Sensitivity of emotion detection from body and physiology components.** Prediction of emotion labels from body and physiological components used separately. Sensitivity rate represents the average of the sensitivity measures for each emotion label across the leave-one-subject-out cross-validation. The error bars show the standard deviation.

Furthermore, since we observed that the body and physiology component was the least effective in discriminating discrete emotions, we also examined the sensitivity rates for each emotion and compared the performance of models using either the body-related CoreGRID items (i.e., subjective ratings), the physiological measures (i.e., objective recordings), or both information (Figure 4B). Consistently, we found that the sensitivity rates obtained with all models using body or physiology information showed a very poor rate of discrimination for the majority of emotions, in comparison to predictions based on other components, except for calm and surprise which were more successfully discriminated. Interestingly, the subjective body-related items from CoreGRID tended to surpass the objective physiology data (*t*(19)= 3.448, *p* = 0.002, Cohen’s *d* = 0.88, 95% *CI* [0.27 1.49]).

## DISCUSSION

The CPM defines emotions by assuming that they are multicomponent phenomena, comprising changes in appraisal, motivation, expression, physiology, and feeling. A considerable advantage of this theory is that it offers the possibility of computational modeling based on a specific parameter space, in order to account for behavioral (Meuleman et al., 2018; Wehrle & Scherer, 2001) and neural (Leitão et al., 2020; Mohammadi et al., 2020) aspects of emotion in terms of dynamic and interactive responses among components. The current research applied multivariate pattern classification analyses for assessing the CPM framework with a range of emotions experienced during movie watching. Through this computational approach, we first investigated the links between physiology manifestations and the four other emotion components proposed by the CPM to determine predictive relationships between them. Second, we investigated whether discrete emotion types can be discriminated from the multicomponent pattern of responses and assessed the importance of each component.

Assuming that physiological responses are intertwined with all components of emotion, we expected that ratings on the 32 CoreGRID features would carry information sufficient to predict corresponding physiological changes. Effectively, SVM classifications provided prediction accuracies significantly better than the chance level. However, information from the CoreGRID items did not allow a high accuracy, even though prediction was simplified by being restricted to a binary distribution. This modest accuracy may be explained by a great variability across participants, which could reduce the generalizability of classifiers when they were applied to all individuals rather than within subject. It might also reflect heterogeneity in intra-individual physiological responses among emotions with similar componential patterns. In parallel, the opposite approach to predict ratings of CoreGRID features based on the means and variances of physiological responses also yielded a performance significantly higher than chance level but still relatively low. Because each CoreGRID item focuses on quite specific behavioral features, it is however not surprising that the sole use of physiology would be insufficient to precisely determine the ratings.

If experiencing an emotion affects simultaneously more than one major component of emotion, one would expect that componential responses are clustered into qualitatively differentiated patterns (Fontaine et al., 2013, Scherer, 2005a). In the CPM view, an emotion arises when components are coherently organized and transiently synchronized (Scherer, 2005b). Accordingly, subjective emotion awareness might emerge as the conscious product of the feeling component generated by such synchronization (Grandjean et al., 2008). However, verbal accounts of conscious feelings may restrict the richness of emotional experience when using only declarative reports. Therefore, we anticipated that integrating the five components together into multivariate pattern analyses would provide higher accuracy rates in emotion prediction. This hypothesis was effectively confirmed, as the best prediction performances were obtained from nonlinear multiclass SVM when the 32 CoreGRID items and the physiological measures were used all together in the model.

On the other hand, the CPM assumes a strong causal link between appraisal and other components of emotion, since appraisal processes are the primary trigger of emotion and should account for a major part of qualitative differences in feelings (Moors & Scherer, 2013). For example, a cross-cultural study demonstrated that an appraisal questionnaire alone (31 appraisal features) could discriminate between 24 emotion terms with an accuracy of 70% (Scherer & Fontaine, 2013). In our study, we found all components provided relevant information and, although being the best predictor, the appraisal component did not provide significantly more information compared to the other components. Conversely, a model using only body and physiology features yielded lower accuracy. Likewise, through our one-component-out model comparisons, we found the body and physiology component was negligible in the overall discrimination of discrete emotion labels when all the other components were present, demonstrating that information derived from the body physiological data was already encoded in other components. Furthermore, in our comparison of models using different components separately, we found no clear superiority of one particular component to discriminate between emotions. Calm stood out as the most recognizable state regardless of the component used. Furthermore, besides fear, love, and satisfaction for which all classifiers were poorly sensitive, at least three components were always predicting one emotion category within the same range of accuracy. Consequently, prediction from components did not significantly differ across emotions, in accordance with a synchronized and combined engagement of these components during emotion elicitation.

In line with our data indicating that physiological measures did not reliably discriminate among emotion categories, the relationship between physiological responses and emotions has long fueled conflicting views. Some authors claimed that there is no invariant and unique autonomic signature linked to each category of emotion (Barrett, 2006), or that physiological response patterns may only distinguish dimensional states (Mauss et al., 2009). In contrast, because emotions imply adaptive and goal-directed reactions, they might trigger differentiated autonomic states to modulate behavior (Kragel & LaBar, 2013; Stemmler, 2004). In this vein, Kreibig (2010) reviewed the most typical ANS responses induced across various emotions and pointed to fairly consistent and stable characteristics for particular affective experiences, but without explicitly confirming a strict emotion specificity since no unique physiological pattern could be highlighted as directly diagnostic of a single emotion. These data suggests that a single or small number of physiological indices are not able to differentiate emotions, but a broader set of measures need to be recorded to obtain discriminative power (Harrison, Kreibig, & Critchley, 2013), including physiology as well as other components. Our results also suggest that the most reliable contribution of physiology could be limited to arousal differentiation, predicting only high or low levels along this dimension. Indeed, our prediction models with physiological measures alone were most accurate in the recognition of calm and surprise labels, two emotions that cannot be defined by valence. Calm describes a passive “neutral” state with an absence of any aroused feeling, while surprise represents a high aroused emotion with undefined or mixed valence. This accords with the view that ANS activation contributes to emotional arousal and reflects more general action dispositions rather than specific emotion types (Schachter & Singer, 1962; Tooby & Cosmides, 1990). Moreover, emotion differentiation afforded by heart and respiration rates mostly concerned arousal changes linked to particular emotions (Kreibig, 2010). More specifically, sympathetic responses were associated with high arousal emotions (anger, fear, joy and surprise), while low arousal emotions (sadness, contentment and affection) were associated with parasympathetic responses (Scherer & Moors, 2019). Altogether, physiology patterning may reflect the type and urgency of action tendencies more than the nature of emotion per se.

Our study is not without limitations. Firstly, statistical machine learning methods may be considered as uninterpretable black boxes. Indeed, SVM analysis gives no explicit clue on functional dimensions underlying classification performances. Secondly, these data-driven methods often need large amounts of data. With our sample of 20 participants, discrimination of specific patterns among the different emotion components was relatively limited. However, we acquired data over a large number of videos and events covering a range of different emotions. Thirdly, although each movie segment was played again before rating each CoreGRID item, participants were asked to answer according to their first feelings during the initial viewing and changes in emotional experience due to repetition or potential recall biases may not be fully ruled out. Fourthly, we used a restricted number of CoreGRID items due to time and experimental constraints. It would certainly be beneficial to measure each component in more detailed ways by taking more features into account. In particular, since the appraisal component is crucial for emotion elicitation, a wider range of appraisal dimensions might allow a more precise discrimination of discrete emotions and physiological patterns. Further, evaluating motor behaviors (e.g. facial expressions) with direct measures such as EMG rather than self-report items could have led to more objective and perhaps more discriminant measures, particularly concerning variations of pleasantness (Larsen, Norris, & Cacioppo, 2003). Lastly, even though using film excerpts has many advantages (e.g., naturalistic and spontaneous emotion elicitation, control over stimuli and timing, standardized validation, and concomitant measurement of physiological responses), an ideal experimental paradigm should evoke first-person emotions in the participants to fully test the assumptions of the CPM framework. In other words, the only way to faithfully elicit a genuine emotion is to get participants to experience an event as pertinent for their own concerns, in order to activate the four most important appraisal features (relevance, implication, coping, normative significance) that are thought to be crucial to trigger an emotion episode (Sander et al., 2005). Viewing film excerpts is an efficient but passive induction technique and, therefore, the meaning of some appraisal components might be ambiguous or difficult to rate. As a result, subjective reports of behaviors and action tendencies were most likely different compared to what they would be for the same event in real life. Future research should develop more ecological scenarios that can be experienced by participants according to their self-relevance and followed by true choices of possible actions. To address that problem, a recent study used virtual reality games to assess the CPM across various emotions (Meuleman & Rudrauf, 2018) and found that fear and joy were predicted by appraisal variables better than by other components, whereas these two emotions were generally poorly classified in our study. Other studies used video game play to assess appraisal and other components during brain imaging (Leitão et al., 2020).

## CONCLUSION

Taken together, our results support the reliability of CPM in the study of emotion. Multivariate pattern classification analyses generated results significantly better than chance level to predict (1) changes in physiological measures from the 32 CoreGRID items, (2) ratings of the majority of CoreGRID items from physiological measures, and (3) discrete emotion labels that refer to conscious feelings experienced by the participants and presumably emerge from a combination of physiological and behavioral parameters. We also found that physiology was interrelated to each of the other major components of emotion, supporting the hypothesis of synchronized recruitment of components during an emotion episode. Moreover, the prediction accuracy for emotion labels was significantly higher when classifiers were trained using the full componential features of emotion. These findings indicate a reliable interindividual consistency of the CPM. However, further work is warranted to determine why certain patterns of behavioral and physiological responses were misclassified into incorrect emotion categories, and study more deeply the links between different emotions. It would be valuable to determine whether poor discrimination stems from a too low sensitivity of our CoreGRID items and physiological measures, or whether some categories of emotions simply cannot be differentiated into distinct entities with such methods, perhaps due to a high degree of overlap within the different components of emotion. Finally, and more generally, further investigations are now also required to understand how distributed mental and bodily changes are integrated into a coherent feeling experience. It is thought that parallel appraisal processes produce an appraisal output representing the affective significance of an event, and concomitantly different outputs are triggered from motivational, physiological, and motor components, but it remains unclear how these representations are then integrated into a feeling component and emerge into awareness. Even if the existence of differentiated autonomic patterning in emotions is confirmed, it would still be necessary to determine whether and to what extent physiological information contributes to subjectively experienced emotional feelings. Further development allowing objective measures for each component during first-person elicitation paradigms are required to limit as much as possible the use of self-assessment questionnaires and ensure ecological validity. Since an emotion is considered as a multicomponent process instigating various cognitive, motivational, physiological, and motor responses simultaneously, the feeling component represents only a small conscious part of this process and understanding the complexity of these relationships may necessitate powerful computational analyses, based on nonlinear multivariate machine learning algorithms and applied to large naturalistic datasets.

## ACKNOWLEDGMENTS

This research was supported by a grant from the Swiss National Science Foundation (SNF Sinergia No. 180319) and the National Centre of Competence in Research (NCCR) Affective Sciences (under grant No. 51NF40-104897) and it was conducted on the imaging platform at the Brain and Behavior Lab (BBL) and benefited from the support of the BBL technical staff.

## SUPPLEMENTAL MATERIAL

**Table S1.** List of the 40 selected film excerpts. Duration (in seconds), the pre-labelled emotions and the number of highly emotional segments are presented.

| Film excerpts | Duration (s) | Related Emotion | Segments |
| --- | --- | --- | --- |
| 12 Years A Slave | 120 | Anger | 4 |
| 28 Days Later | 143 | Fear | 4 |
| 28 Days Later | 49 | Anxiety | 4 |
| 28 Days Later | 125 | Fear | 4 |
| A Fish Called Wanda | 173 | Joy | 4 |
| American History X | 77 | Anxiety | 4 |
| Bambi | 86 | Love | 2 |
| Batman Returns | 73 | Surprise | 3 |
| Cry Freedom | 132 | Disgust | 4 |
| Dangerous Minds | 128 | Sadness | 2 |
| Edward Scissorhands | 93 | Calm | 4 |
| Forrest Gump | 121 | Love | 3 |
| Ghost | 215 | Love | 4 |
| Hotel Rwanda | 209 | Sadness | 4 |
| Hotel Rwanda | 96 | Sadness | 2 |
| Hotel Rwanda | 97 | Anger | 3 |
| Kill Bill 1 | 68 | Surprise | 2 |
| Kill Bill 1 | 144 | Surprise | 2 |
| Life Is Beautiful | 127 | Anger | 3 |
| Life Is Beautiful | 105 | Love | 2 |
| Love Actually | 76 | Joy | 3 |
| Love Actually | 104 | Joy | 4 |
| Love Actually | 28 | Satisfaction | 1 |
| Mr. Bean's Holiday | 128 | Joy | 4 |
| My Girl | 146 | Sadness | 2 |
| Planet Earth | 66 | Calm | 3 |
| Pride And Prejudice | 91 | Calm | 3 |
| Remember The Titans | 132 | Satisfaction | 3 |
| Schindler's List | 115 | Disgust | 2 |
| Scream 2 | 215 | Fear | 4 |
| Terminator 2 | 131 | Satisfaction | 4 |
| The Dentist | 56 | Fear | 2 |
| The Fly | 81 | Anxiety | 3 |
| The Pianist | 81 | Anger | 2 |
| The Pianist | 117 | Anxiety | 2 |
| There Is Something About Mary | 175 | Surprise | 4 |
| The Shining | 85 | Satisfaction | 2 |
| Trainspotting | 104 | Disgust | 3 |
| Trainspotting | 62 | Disgust | 1 |
| Wild Alaska | 62 | Calm | 2 |

**Figure S1.**
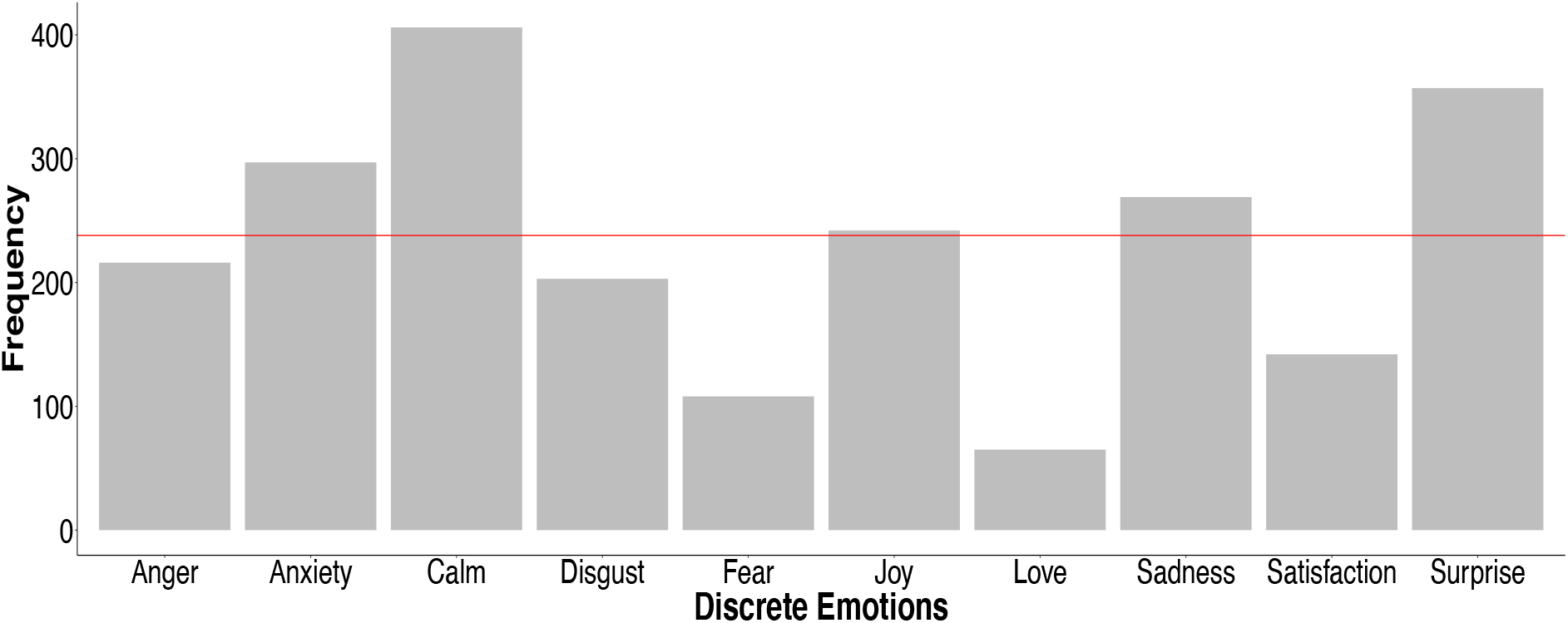
Histogram of discrete emotions. Histogram of categorical emotions based on their frequency in the assessments of 119 emotional events per 20 participants. The red line indicates the ideal frequency if the samples were equally distributed.

